# Analysis of an HIV model with post-treatment control

**DOI:** 10.1101/448308

**Authors:** Shaoli Wang, Fei Xu

**Affiliations:** School of Mathematics and Statistics, Henan University, Kaifeng 475001, Henan, PR China; Department of Mathematics, Wilfrid Laurier University, Waterloo, Ontario, Canada N2L 3C5

## Abstract

Recent investigation indicated that latent reservoir and immune impairment are responsible for the post-treatment control of HIV infection. In this paper, we simplify the disease model with latent reservoir and immune impairment and perform a series of mathematical analysis. We obtain the basic infection reproductive number *R*_0_ to characterize the viral dynamics. We prove that when *R*_0_ < 1, the uninfected equilibrium of the proposed model is globally asymptotically stable. When *R*_0_ > 1, we obtain two thresholds, the post-treatment immune control threshold and the elite control threshold. The model has bistable behaviors in the interval between the two thresholds. If the proliferation rate of CTLs is less than the post-treatment immune control threshold, the model does not have positive equilibria. In this case, the immune free equilibrium is stable and the system will have virus rebound. On the other hand, when the proliferation rate of CTLs is greater than the elite control threshold, the system has stable positive immune equilibrium and unstable immune free equilibrium. Thus, the system is under elite control.

**Author summary:** In this article, we use mathematical model to investigate the combined effect of latent reservoir and immune impairment on the post-treatment control of HIV infection. By simplifying an HIV model with latent reservoir and immune impairment, and performing mathematical analysis, we obtain the post-treatment immune control threshold and the elite control threshold for the HIV dynamics when *R*_0_ > 1. The HIV model displays bistable behaviors in the interval between the two thresholds. We illustrate our results using both mathematical analysis and numerical simulation. Our result is consistent with recent medical experiment. We show that patient with low proliferation rate of CTLs may undergo virus rebound, and patient with high proliferation rate of CTLs may obtain elite control of HIV infection. We perform bifurcation analysis to illustrate the infection status of patient with the variation of proliferation rate of CTLs, which potentially explain the reason behind different outcomes among HIV patients.

---

Introduction In 2010, an HIV-infected mother gave birth to a baby prematurely in a Mississippi clinic. The infant was known as the ‘Mississippi baby’. Before delivery, the mother was not diagnosed with HIV infection did not receive antiretroviral treatment [26]. At the age of 30 hours, the baby received liquid, triple-drug antiretroviral treatment. Such treatment was terminated at the age of 18 months and since then, the virus level in the baby remains undetectable. Though it was thought that the baby was cured of HIV, a routine clinical test on July 10, 2014 showed that the level of virus in the ‘Mississippi baby’ became detectable (16,750 copies/ml) [26].

Antiretroviral therapy (ART) is effective in inhibiting the HIV infection and prolongs the life of infected individuals. However, due to the existence of latent reservoirs, it is unable to totally eliminate the virus infection[7, 8, 12, 13, 48]. The time it takes the virus to rebound varies. For example, the virus level of the Mississippi baby remains undetectable for years before the virus rebound [26, 30]. Sometimes, a host may have low virus load after antiretroviral therapy. Investigations have been carried out to reveal the causes of low virus level and virus rebound[9, 30, 38].

Conway and Perelson constructed a mathematical model to investigate the dynamics of virus rebound [9]. Their investigation reveals the interplay between immune response and latent reservoir, and shows that post-treatment control may appear. Recent investigations indicated that early antiretroviral therapy may be responsible for the development of post-treatment control with plasma virus remaining undetectable after the cessation of treatment. However, only a small proportion of patients receiving early antiretroviral therapy developed post-treatment control. Further investigations are to be carried out to reveal the reasons behind this.

Treasure et al investigated the HIV rebound in patients who terminated the antiretroviral therapy. They showed that a patient who discontinued the antiretroviral therapy may or may not undergo immediate HIV rebound[38].

As an important approach to investigate disease transmission, mathematical modeling provides insights into interactions between viral and host factors. Evaluating the behaviors of the viral models yields a better understanding of the disease and is beneficial to the development of appropriate therapy strategies. In the literature, mathematical models of within-host viral dynamics have been designed [1, 3, 10, 11, 15, 27–29, 44–46]. Immune response has also been integrated into within host models to investigate the combined effects of viral dynamics and immune process of the host [6, 16, 23, 36, 40, 41, 43, 49].

Regoes et al. [32] incorporated immune impairment into viral models to consider the effects that target cell limitation and immune responses have on the evolution of virus. Their investigations indicated that the immune system of the host may collapse when the impairment rate of HIV surpasses its threshold value. Iwami et al. [17, 18] investigated the HIV dynamics with immune impairment using mathematical models. The authors got the ‘risky threshold’ and ‘immunodeficiency threshold’ by performing analysis. The results implied that the immune system always collapses when the impairment rate is greater than the threshold value. Immune impairment in within-host virus models have received much attention in the literature [2, 37, 39].

HIV latent reservoir is responsible for the rebound in HIV viral load. As a major barrier to the eradication of HIV-1 virus, latent reservoir poses persistent risks to the hosts. The infected cells in the latent reservoir remain undetectable to the immune system and can be reactivated to produce virions with the termination of drug therapy [19, 20, 33, 34, 42]. Investigations showed that the size of the virus reservoir is relatively stable [42]. For a patient under sufficient antiretroviral therapy (ART), ongoing viral replication rate in the reservoir remains low [19]. However, for infected individuals under ART of lower efficiency, there might be coexistence of latent reservoir and virus. Rong and Perelson [34] performed a thorough study on the replenishment of the latent reservoir induced by latently infected cells that are occasionally reactivated. The authors indicated that such scenario corresponds to the half-life of the latent reservoir.

Post-treatment control of HIV attracted the attention of researchers. Conway and Perelson integrated the post treatment into an HIV model and performed analysis [9]. Here, we simplify the model proposed in [9] to obtain

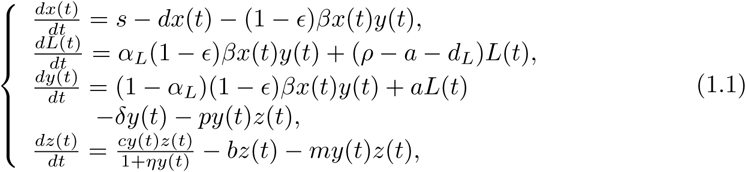

where *x* denotes the concentration of activated CD4^+^ T cells, *L* latently infected cells, *y* productively infected CD4^+^ T cells and *z* the immune cells. The effectiveness of both drug classes is represented by ϵ ∈ [0, 1]. Here *ϵ* is also known as the overall treatment effectiveness of HIV. If the treatment is terminated, *ϵ* = 0. If the therapy is 100% effective, we have *ϵ* = 1 [9, 33].

In the literature, the immune and immune impairment function 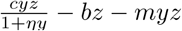 has been applied to the viral models to characterize the interaction between the immune cells and the productively infected CD4^+^ T cells [11, 31, 39]. Wang and Liu [39] constructed a within-host viral dynamics models to consider HIV infection with immune impairment. In this article, we consider the post-treatment immune control, the biological implication behind the ‘Mississippi baby’. By mathematical analysis, we obtain the threshold of proliferation rate of CTLs, which determines the HIV infection status. We also perform bifurcation analysis and demonstrate the bistable behavior of the model, which is consistence with results from recent medical trial.

## 1 Preparation

In this section, we perform mathematical analysis for the model (1.1). We prove the positiveness and boundedness of the solutions to system (1.1) and calculate the equilibria of the model.

### 1.1 Positiveness and boundedness

In the following, we show that system (1.1) is well-posed.

#### Theorem 2.1.

System (1.1) has a unique nonnegative solution with initial values 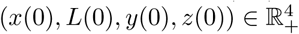, where 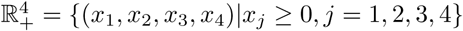.

Furthermore, the solution is bounded.

**Proof.** It follows from the fundamental theory of ordinary differential equations [14] that there exists a unique solution to system (1.1) with nonnegative initial conditions.

For any nonnegative initial data, let *t*_1_ > 0 be the first time when *x*(*t*_1_) = 0. From the first equation of (1.1) we have that 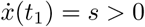, which implies that *x*(*t*) < 0 for *t* ∈ (*t*_1_ − ∊_1_, *t*_1_), where *∊*_1_ is an arbitrarily small positive constant. This is a contradiction. Therefore, *x*(*t*) is always positive. Since *z* = 0 is a constant solution of the last equation of (1.1), it follows from the fundamental existence and uniqueness theorem that *z* > 0 for all *t* > 0.

Suppose there is a first time *t*_2_ > 0 when *y*(*t*_2_)*z*(*t*_2_) = 0. Then we have

i. *L*(*t*_2_) = 0, *y*(*t*) ≥ 0 for *t* ∈ [0, *t*_2_], or
ii. *y*(*t*_2_) = 0, *L*(*t*) ≥ 0 for *t* ∈ [0, *t*_2_].

For case(i), since *x*(*t*) is positive, it follows from the variation of constants formula that 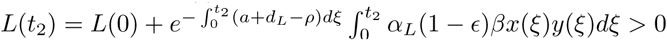, which is in contradiction to *L*(*t*_2_) = 0.

For case (ii), the third equation of system (1.1) implies that 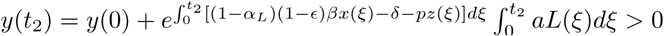, which is in contradiction to *y*(*t*_2_) = 0. Thus, *L*(*t*) and *y*(*t*) are always positive.

Next, we expatiate upon the boundedness of solutions of (1.1). Let

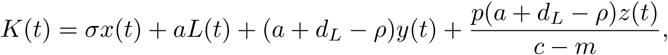

where *σ* = aα_*L*_ + (1 − α_*L*_)(*a* + *d_L_* − ρ). Since all solutions of (1.1) are positive, we have

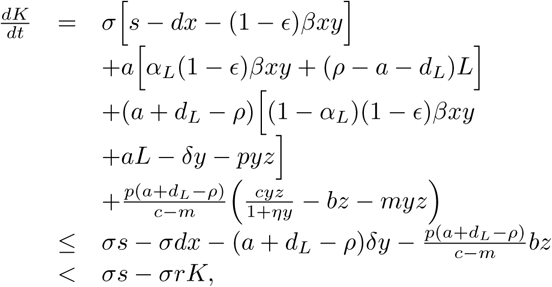

where 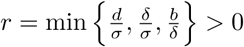. Let φ denote the solution to the following system

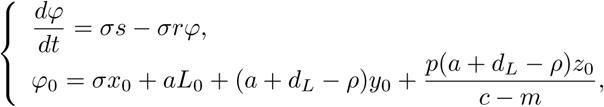

where *x*_0_, *y*_0_ and *z*_0_ are the initial values of system (1.1) and φ_0_ = *K*_0_ > 0. We then obtain 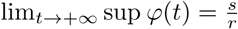. By comparison theorem [35], we get *K*(*t*) < φ(*t*). Therefore, *x*(*t*), *L*(*t*), *y*(*t*) and *z*(*t*) are bounded.

### 1.2 Equilibria

In this section, we consider the existence of the equilibria to system (1.1).

i. If *R*_0_ < 1, system (1.1) only has an infection-free equilibrium 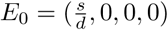, where

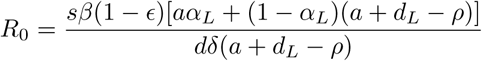

is the basic infection reproductive number. Here, *R*_0_ is the expected number of newly infected cells generated from an infected cell at the beginning of the infectious process.
ii. If *R*_0_ > 1, system (1.1) also has an immune-free equilibrium *E*_1_ = (*x*_1_, *L*_1_, *y*_1_, 0), where

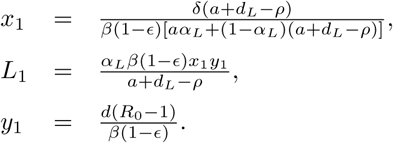 Solving equation 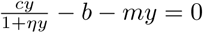 yields two positive roots, given by 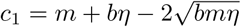 and 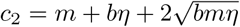. We then get the existence conditions for the positive equilibria.
iii. If 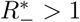 and *c* > *c*_2_, system (1.1) has an immune equilibrium 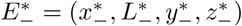. If 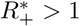 and *c* > *c*_2_, system (1.1) has an immune equilibrium 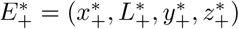 as well. Here

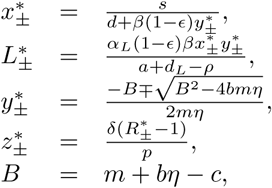

and

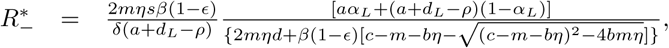 Denote

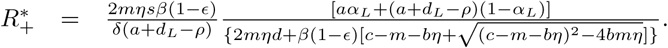

and

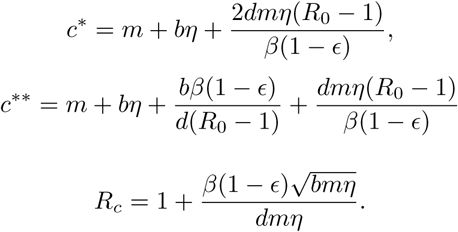

We then have the following results.

#### Lemma 1.1.

*R*_0_ > *R_c_* > 1, ⇔ *c*\* > *c*\*.

**Proof.**

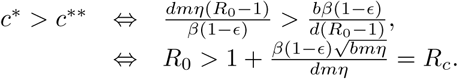

#### Lemma 1.2.

(i) *R*_0_ > *R*_c_ > 1, *c*\* > *c*_2_. (ii) 1 < *R*_0_ < *R*_c_ ⇔ *c*\* < *c*_2_.

**Proof.**

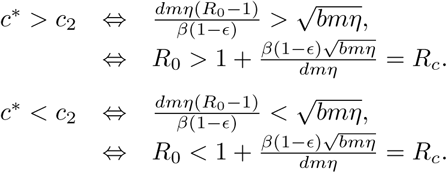

#### Lemma 1.3.

(i) Assume 1 < *R*_0_ < *R_c_*. If 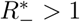, then *c* > *c*\*\*. (ii) Assume *R*_0_ > *R_c_* > 1. If 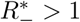, then *c* > *c*_2_.

**Proof.**

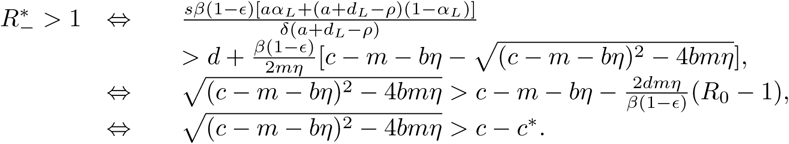

If *c* < *c*\* and one of the conditions *c* < *c*_1_ or *c* > *c*_2_ holds, then 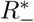 is always greater than one. If *c* > *c*\*, solving 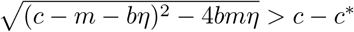 yields *c* > *c*\*.

i. If 1 < *R*_0_ < *R_c_*, then *c*\* < *c*_2_. From 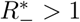, we can deduce that *c* > *c*\*.
ii. If *R*_0_ > *R_c_* > 1, then *c*\* > *c*_2_. From 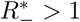, we can deduce that *c* > *c*_2_.

#### Lemma 1.4.

(i) If 1 < *R*_0_ < *R_c_*, then 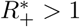 has no solution. (ii) Assume that *R*_0_ > *R_c_* > 1. If 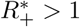, then *c*_2_ < *c* < *c*.

**Proof.**

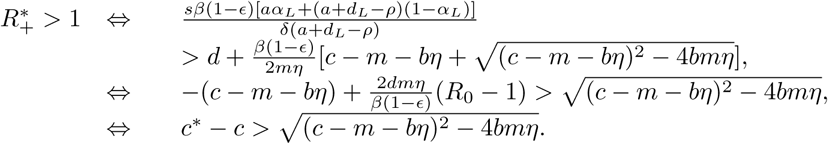

(i) If 1 < *R*_0_ < *R_c_*, then *c*\* < *c*_2_. Thus 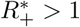 has no solution. (ii) If *R*_0_ > *R_c_* > 1, then *c*\* > *c*_2_. Solving 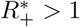, we have *c*_2_ < *c* < *c*\*\*.

By Lemma 2.1∼2.4, summing up the above analysis yields the existence results of the equilibria of system (1.1)

#### Theorem 1.2

i. System (1.1) always has an infection-free equilibrium *E*_0_.
ii. If *R*_0_ > 1, system (1.1) also has an immune-free equilibrium *E*_1_.
iii. If 1 < *R*_0_ < *R_c_* and *c* > *c*\*\*, system (1.1) has only one positive equilibrium 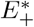.
iv. If *R*_0_ > *R_c_* > 1 and *c*_2_ < *c* < *c*\*\*, system (1.1) has two positive equilibria 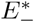 and 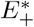. While *R*_0_ > *R_c_* and *c* > *c*\*\*, system (1.1) has only one positive equilibrium 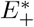. The existence of the positive equilibria of the model is summarized in Tables 1 and 2.

**Table 1.**
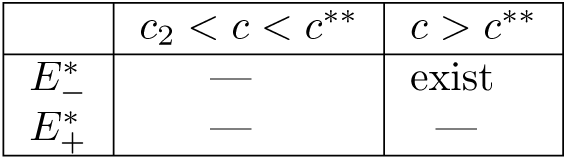
The existence of the positive equilibria when 1 < *R*_0_ < *R_c_*.

**Table 2.** The existence of the positive equilibria when *R*_0_ > *R_c_* > 1.

| | $c_2 < c < c^{**}$ | $c > c^{**}$ |
| --- | --- | --- |
| $E_-^*$ | exist | exist |
| $E_+^*$ | exist | — |

## 2 Stability analysis

In this section, we consider the stability of the equilibria of system (1.1).

Let 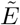 be any arbitrary equilibrium of system (1.1). Its corresponding Jacobian matrix is obtained as

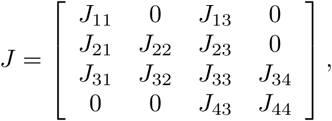

where

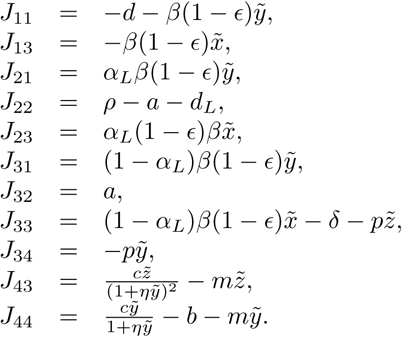

The characteristic equation of the linearized system of (1.1) at 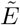 is given by

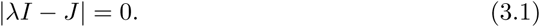

### 2.1 Stability analysis of Equilibrium *E*_0_

#### Theorem 2.1.

If *R*_0_ < 1, then the infection-free equilibrium *E*_0_ of system (1.1) is locally asymptotically stable. If *R*_0_ > 1, then *E*_0_ is unstable.

**Proof.** For equilibrium *E*_0_(*x*_0_, 0, 0, 0), the characteristic equation (3.1) reduces to

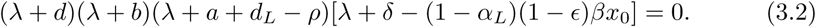

It is easy to see that equation (3.2) has two negative roots, obtained as

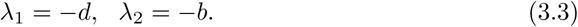

The other eigenvalues are determined by

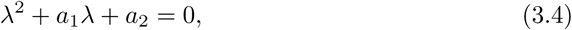

where

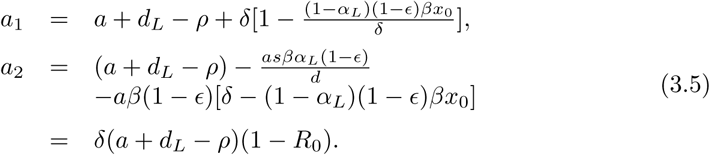

If *R*_0_ < 1, we have *a*_1_ > 0 and *a*_2_ > 0, and as such equation (3.4) has two negative roots. Thus, *E*_0_ is locally stable for *R*_0_ < 1.

If *R*_0_ > 1, from (3.5) we know that *E*_0_ is a saddle, and hence unstable. The proof of Theorem 3.1 is complete.

#### Theorem 2.2.

If *R*_0_ < 1, then the infection-free equilibrium *E*_0_ of system (1.1) is globally asymptotically stable.

**Proof.** Define a function

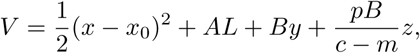

where *A* and *B* are undetermined positive coefficients. It is easy to see that *V* is a positive Lyapunov function. Evaluating the time derivative of *V* along the solution of system (1.1) yields

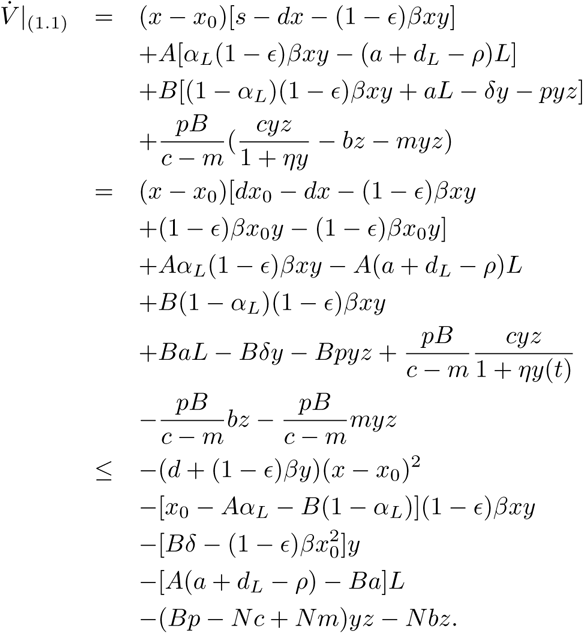

If we choose

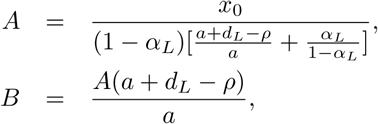

then

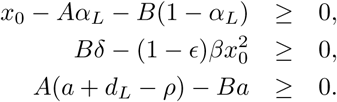

Thus, if *R*_0_ ≤ 1, then 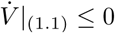. Since *x, L, y, z* are positive, we have 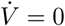 if and only if (*x, L, y, z*) = (*x*_0_, 0, 0, 0). Therefore, it follows from the classical Krasovskii-LaSalle principle [21, 22] that *E*_0_ is globally asymptotically stable.

Biologically, the global asymptotic stability of the uninfected equilibrium *E*_0_ of system (1.1) implies that the virus will die out in the host if the treatment is strong enough to ensure *R*_0_ < 1.

### 2.2 Stability analysis of Equilibrium *E*_1_

Now we consider the stability of equilibrium *E*_1_.

#### Theorem 3.3.

Suppose that the immune-free equilibrium exists (i.e., *R*_0_ > 1). When 0 < *c* < *c*\*\*, *E*_1_ is locally asymptotically stable. When *c* > *c*\*\*, *E*_1_ is unstable.

**Proof.** The characteristic equation of the linearized system of (1.1) at *E*_1_ is given by

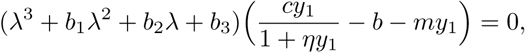

where

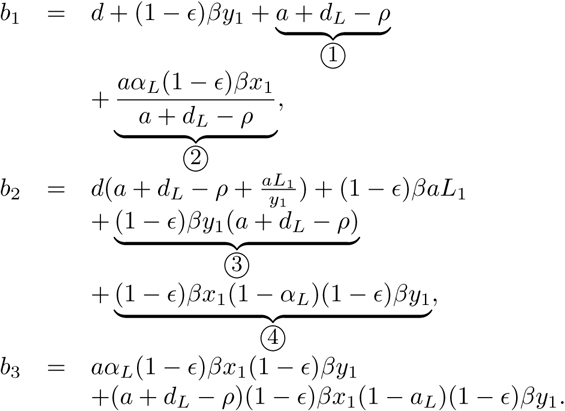

Clearly,

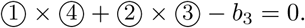

Thus, we have *b*_1_*b*_2_ − *b*_3_ > 0. We then consider the sign of the eigenvalue

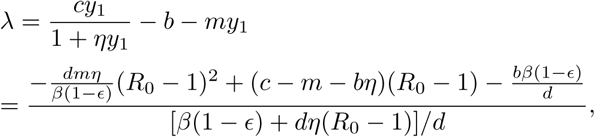

which is determined by

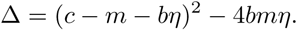

Let Δ = 0, we have *c* = *c*_1_ or *c* = *c*_2_.

i. If = 0, then *c* = *c*_1_ or *c* = *c*_2_, which is a critical situation.
ii. If Δ < 0, then *c*_1_ < *c* < *c*_2_, and we have λ < 0.
iii. If Δ > 0, then *c* < *c*_1_ or *c* > *c*_2_. To get λ < 0, we must ensure *c* < *m* + *bη* and *R*_0_ < 1 + *R*_1_, or *R*_0_ > 1 + *R*_2_. Meanwhile, from *R*_0_ < 1 + *R*_1_ and *R*_0_ > 1 + *R*_2_, we have *c* < *c*\*\*. Here 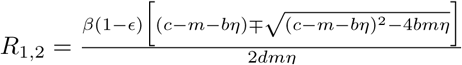. In view of *c*_2_ < *c*\*\*, if *c* < *m* + *bη* or *c*_2_ < *c* < *c*\*\*, then the eigenvalue λ < 0. If *c* > *c*\*\*, we have λ > 0.

In summary, if *c* < *c*_2_ or *c*_2_ < *c* < *c*\*\*, then λ < 0. By the Routh-Hurartz criterion, for *R*_0_ > 1, if *c* < *c*_2_ or *c*_2_ < *c* < *c*\*\*, the equilibrium *E*_1_ of system (1.1) is locally asymptotically stable. If *c* > *c*\*\*, *E*_1_ is unstable.

Biologically, if the proliferation rate of CTLs is less than the critical value *c*, the viral load can be at high level.

### 2.3 Stability analysis of positive equilibria

In this subsection, we consider the stability of the positive equilibria. Here, we use *E*\* = (*x*, L*, y*, z\**) to denote a positive equilibrium of system (1.1).

#### Theorem 3.4.

i. Assume 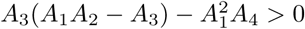. If

(**A.1**) 1 < *R*_0_ < *R_c_* and *c* > *c*\*\*, or
(**A.2**) *R*_0_ > *R_c_* > 1 and *c* > *c*_2_,

system (1.1) has an immune equilibrium 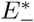, which is a stable node.
ii. If *R*_0_ > *R_c_* > 1 and *c*_2_ < *c* < *c*\*\*, system (1.1) also has an immune equilibrium 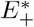, which is an unstable saddle.

**Proof.** The characteristic equation of the linearized system of (1.1) at an arbitrary positive equilibrium *E*\* is given by

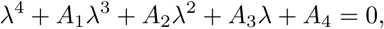

where

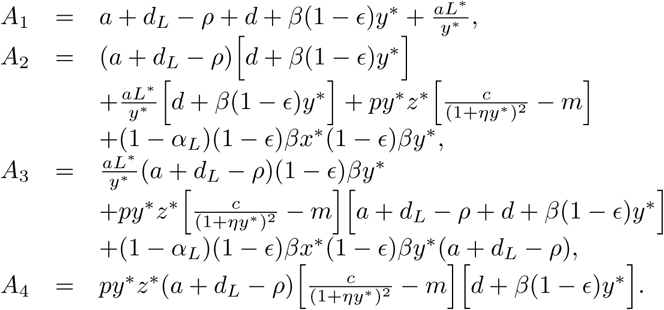

Then we have

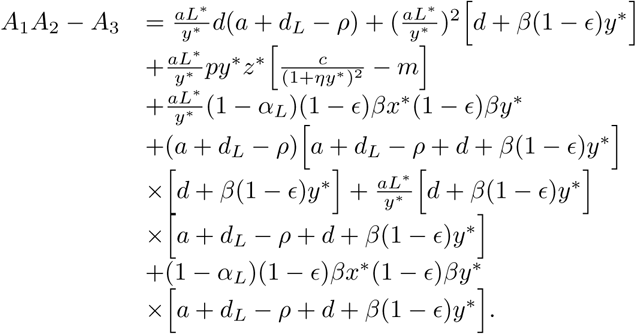

i. For equilibrium 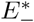, if *c* > *c*_2_, we have 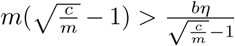. It thus follows that 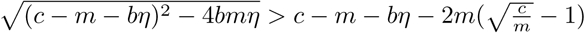. Therefore, 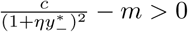. Clearly, *A_i_* > 0, *i* = 1, 2, 3 and *A*_1_*A*_2_ − *A*_3_ > 0. If 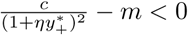, by Routh-Hurartz Criterion, we know that the positive equilibrium 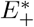 is a stable node in this case.
ii. For equilibrium 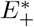, if *R*_0_ > *R_c_* > 1 and *c*_2_ < *c* < *c*\*\*, then 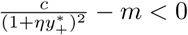 and *A*_4_ < 0. Thus, equilibrium 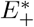 is an unstable saddle for *R*_0_ > *R_c_* and *c*_2_ < *c* < *c*\*\*.

By Theorem 3.3 and Theorem 3.4, we have the following result.

#### Theorem 3.5.

If *R*_0_ > *R_c_* > 1 and *c* = *c*_2_, the immune equilibrium 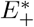 and 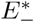 coincide with each other and a saddle-node bifurcation occurs when *c* passes through *c*_2_.

The stabilities of the equilibria and the behaviors of system (1.1) are summarized in Tables 3 and 4.

**Table 3.** The stabilities of the equilibria and the behaviors of system (1.1) in the case 1 < *R*_0_ < *R_c_*. Here, *c*\*\* is the critical value, and we assume 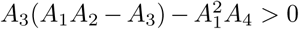.

| | $E_0$ | $E_1$ | $E_-^*$ | $E_+^*$ | System (1.1) |
| --- | --- | --- | --- | --- | --- |
| $R_0 < 1$ | GAS | — | — | — | Converges to $E_0$ |
| $1 < R_0 < R_c, 0 < c < c^{**}$ | US | LAS | — | — | Converges to $E_1$ |
| $1 < R_0 < R_c, c^{**} < c$ | US | US | LAS | — | Converges to $E_+^*$ |

**Table 4.** The stabilities of the equilibria and the behaviors of system (1.1) in the case *R*_0_ > *R_c_* > 1. Here, *c*_2_, *c*\* and *c*\*\* are critical values, and *c*_2_ is a saddle-node bifurcation point. Here we assume 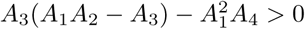.

| | $E_0$ | $E_1$ | $E_-^*$ | $E_+^*$ | System (1.1) |
| --- | --- | --- | --- | --- | --- |
| $R_0 < 1$ | GAS | — | — | — | Converges to $E_0$ |
| $R_0 > 1, 0 < c < c_2$, | US | LAS | — | — | Converges to $E_1$ |
| $R_0 > R_c > 1, c_2 < c < c^{**}$ | US | LAS | LAS | US | Bistable |
| $R_0 > R_c > 1, c^{**} < c < c^*$ | US | US | LAS | US | Converges to $E_+^*$ |
| $R_0 > R_c > 1, c > c^{**}$ | US | US | LAS | — | Converges to $E_+^*$ |

## 3 Sensitive analysis and numerical simulations

### 3.1 Sensitive analysis

Sensitive analysis provides insights into the basic infection reproductive number *R*_0_ with respect to system parameters [47]. In this section, we use latin hypercube sampling (LHS) and partial rank correlation coefficients (PRCCs) [4, 24] to reveal the dependence of the basic infection reproduction number *R*_0_ on a variety of system parameters. As a statistical sampling method, LHS provides efficient analysis of parameter variations across simultaneous uncertainty ranges in each parameter [4]. PRCC, which is obtained from the rank transformed LHS matrix and output matrix [24], indicates the parameters that have the most significant influences on the behaviors of the model. In this work, we perform 4000 simulations per run. We use a uniform distribution function to test the PRCCs for a variety of system parameters.

The PRCC results of the model, Fig. 1, illustrate the dependence of *R*_0_ on different system parameters. The estimations of the distributions for *R*_0_ is approximately a normal distribution. We use |PRCC| as an index to test if the parameter has important correlation with the infection reproduction number *R*_0_. If |PRCC| > 0.4, we say that the correlation is strong. If 0.4 ≥ |PRCC| > 0.2, we say that the correlation is moderate. For 0.2 ≥ |PRCC| > 0, there correlation is weak. As is shown in Fig. 1, the general rate of CD4^+^ T cells *s*, the decay rate of CD4^+^ T cells *d*, the infection rate of CD4^+^ T cells β, the drug efficacy *ϵ* and the latently infected cell death rate *d_L_* have significant influence on the infection reproduction number *R*_0_.

**Fig 1.**
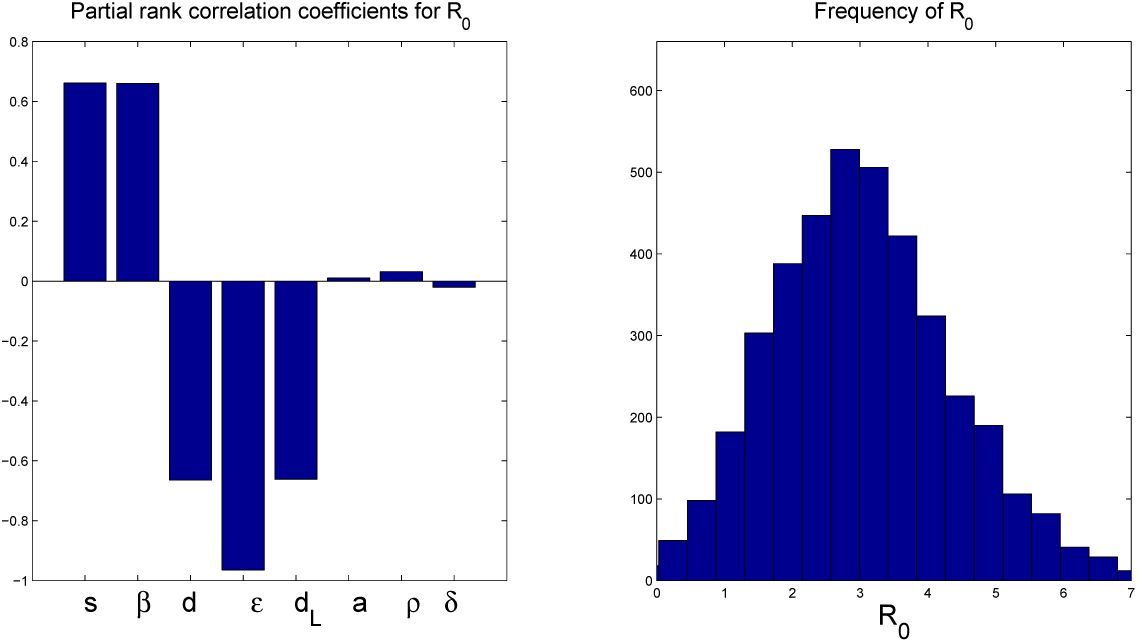
Partial rank correlation coefficients for *R*_0_ and the frequency distribution of *R*_0_. The parameters are shown in Table 5.

### 3.2 Numerical simulations

In this section, we carry out numerical simulations to consider the HIV dynamics of our model. The parameter values are listed in Table 5. We then calculate the thresholds *R*_0_ ≈ 3.0030 > 1, *R_c_* ≈ 1.4243, *c*_2_ ≈ 0.2914 and *c*\*\* ≈ 0.4988. Notice that 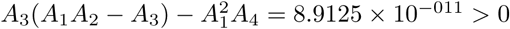. We then get the bistable interval (0.2914, 0.4988). In this case, when *c* < *c*_2_, the immune-free equilibrium *E*_1_ is stable. When *c*_2_ < *c* < *c*\*\*, the immune-free equilibrium *E*_1_ and the positive equilibrium 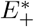 are stable. When *c* > *c*\*\*, only the positive equilibrium 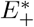 is stable.

**Table 5.** Parameters for the model.

| Symbol | Description | Value | Reference |
| --- | --- | --- | --- |
| $s$ | Proliferation rate of $CD4^+$ T cells | 10 cells / $\mu$ L / day | [5] |
| $d$ | Decay rate of $CD4^+$ T cells | $0.01 \text{ day}^{-1}$ | [5] |
| $\beta$ | Infection rate of $CD4^+$ T cells | $0.015 \mu \text{ L} / \text{day}$ | — |
| $\epsilon$ | Drug efficacy | 0.8 | — |
| $\alpha_L$ | Fraction of newly infected cells that become latently infected | 0.001 | — |
| $\rho$ | Proliferation rate of latently infected cells | $0.0045 \text{ day}^{-1}$ | [9] |
| $a$ | Activation rate | $0.001 \text{ day}^{-1}$ | [9] |
| $d_L$ | Latently infected cell death rate | $0.004 \text{ day}^{-1}$ | [9] |
| $\delta$ | Infected cell death rate | $1 \text{ day}^{-1}$ | [25] |
| $p$ | Killing rate of infected $CD4^+$ T cells | $0.42 \text{ day}^{-1}$ | — |
| $c$ | Proliferation rate of CTLs | $0.45 \text{ day}^{-1}$ | — |
| $\eta$ | Effector cell production Hill function scaling | 1 cells/$\mu$ L | — |
| $b$ | Decay rate of CTLs | $0.1 \text{ day}^{-1}$ | — |
| $m$ | Immune impairment rate of viral | $0.05 \text{ cells} / \mu \text{ L} / \text{day}$ | — |

Fig. 2 shows that there is no positive equilibrium if *c* < 0.2914 and a saddle-node bifurcation appear when *c* passes through 0.2914. The system display bistable behavior for 0.2914 < *c* < 0.4988. As an example, we simulate the time history of the system for *c* = 0.45 ∈ (0.2914, 0.4988) with different initial conditions (see Fig. 3). We find that, with the same parameter values and different initial conditions, the system may converge to different equilibriums. Such simulation result is consistent with recent clinic trial performed by Treasure et al [38].

**Fig 2.**
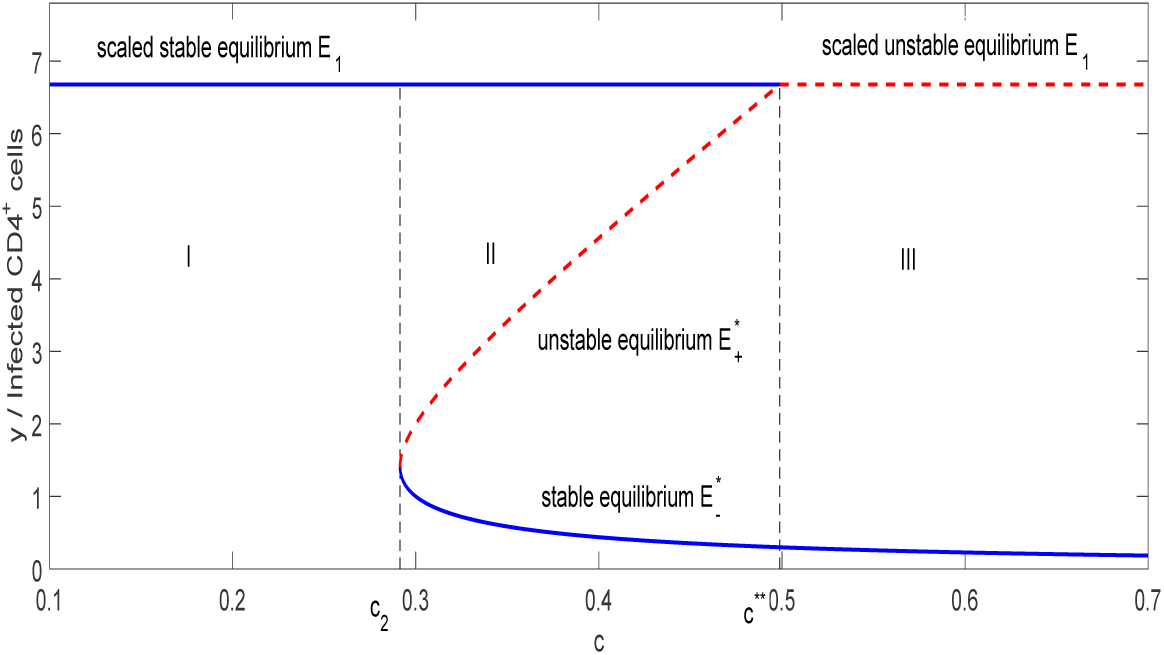
Bistability and saddle-node bifurcation diagram of system (1.1). Here *c* = 0.2914 is a saddle-node bifurcation (SN) point. The bistable interval is (0.2914, 0.4988). The parameter values are shown in Table 5. There are three phases in this figure. In phase I (0 < *c* < *c*_2_), the system has virus rebound. In phase II (*c*_2_ < *c* < *c*\*\*), the system has bistable behavior. In phase III (*c* > *c*\*\*), the system is under elite control.

**Fig 3.**
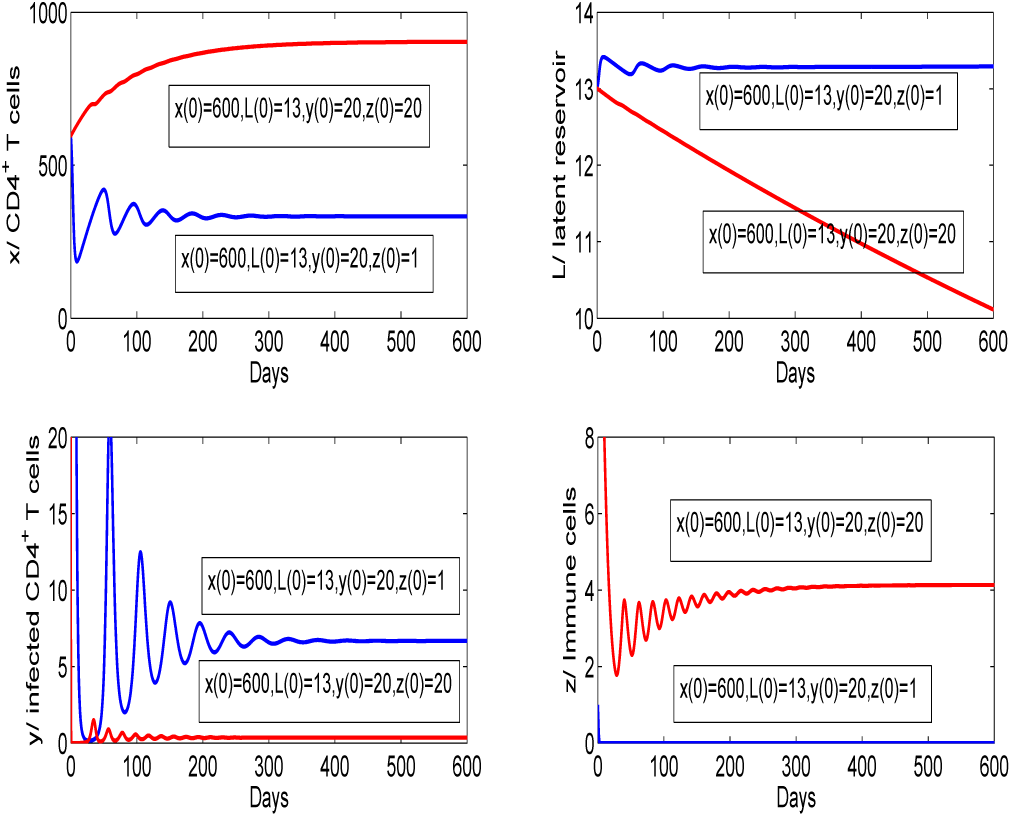
Time history of system (1.1) for *c* = 0.45 (*c*_2_ < *c* < *c*\*\*). All the other parameter values are listed in Table 5. The trajectories of system (1.1) converge to different equilibria for different initial values, i.e., system (1.1) has bistable behavior. The initial values are *x* (0) = 600, *L* (0) = 13, *y* (0) = 20, *z*(0) = 1 (blue) and *x* (0) = 600, *L* (0) = 13, *y* (0) = 20, *z* (0) = 20 (red).

We also consider the influence of system parameters on the elite control threshold *c*\*\* by PRCCs. Fig. 4 shows that the immune impairment rate of virus *m* and the proliferation rate of latently infected cells *ρ* are positively correlated with the elite control threshold *c*\*\*. On the other hand, the death rate of infected cells δ has negative correlation with the elite control threshold *c*\*\*. It thus follows that decreasing immune impairment rate *m* is beneficial for obtaining post-treatment immune control. Decrease the immune impairment rate *m* and the proliferation rate of latently infected cells *ρ*, and increasing the death rate of infected cells δ are beneficial for the host to get elite control.

**Fig 4.**
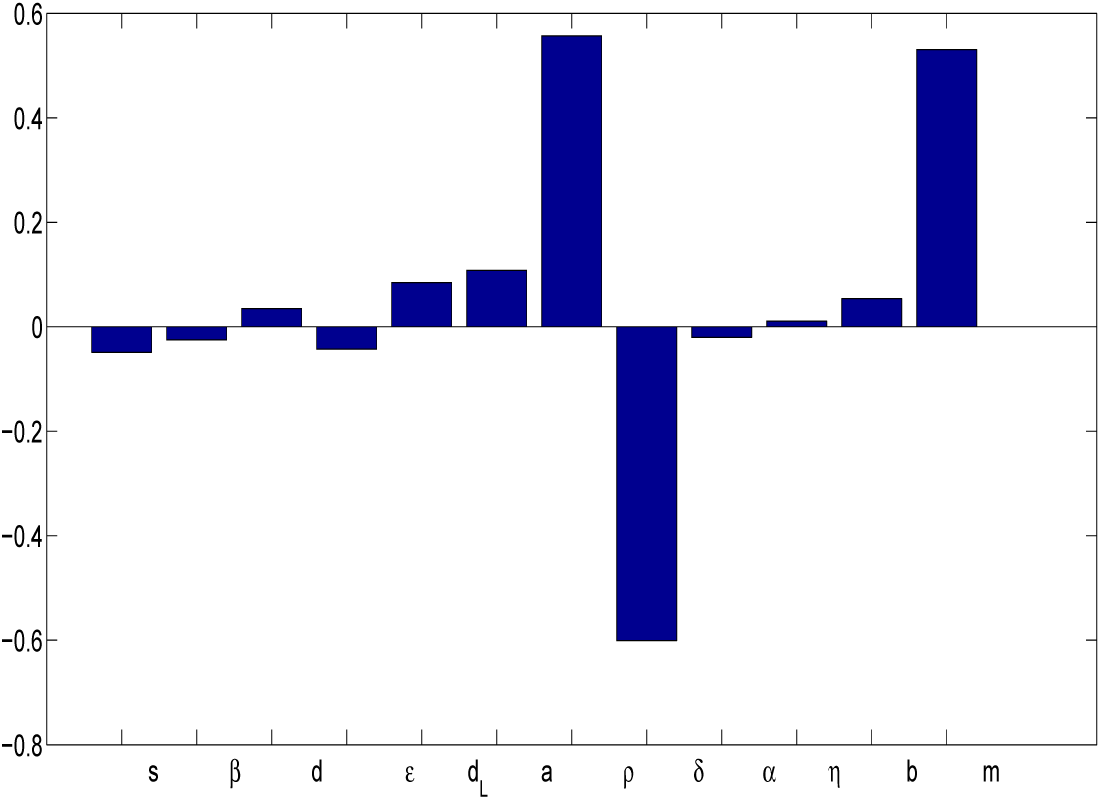
Partial rank correlation coefficients for *c*\*\*. The parameter values are shown in Table 5.

## 4 Discussion

In this paper, we investigate the viral dynamics of a simplified within host model. By performing mathematical analysis and numerical simulations, we obtain the post-treatment immune control threshold and the elite control threshold. We get conditions for the model to reach post-treatment immune control and elite control.

The expression of the post treatment control threshold implies that the immune impairment rate of virus *m* has positive correlation with the post treatment control threshold. Early initiation of ART after infection allows PTC by limiting the size of latent reservoir. A patient with latent HIV reservoir small enough may obtain adaptive immune response to prevent viral rebound (VR), and thus has controlled infection Conway and Perelson [9].

Sensitive analysis and numerical simulations imply that decreasing the immune impairment rate is beneficial for the host obtain post-treatment immune control and the elite control. A comprehensive HIV treatment involving decreasing the immune impairment rate of virus, decay rate of CTLs and effector cell production Hill function scaling allows the host to obtain elite control efficiently.

The proliferation rate of latently infected cells *ρ* plays an important role in the elite control. It is worth carrying out further investigation to reveal the viral dynamics of the within host model with logistic proliferation rate of latently infected cells, given by system (5.1).

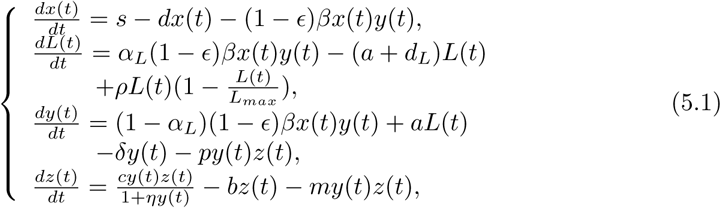

Using the same method of analyzing system (1.1), we can get theoretical results. Here, we carry out numerical simulations to show its bistable behaviors. As shown in Fig.5, if we choose parameters listed in Table 5 and *L_max_* = 50, system (5.1) displays bistable behaviors.

**Fig 5.**
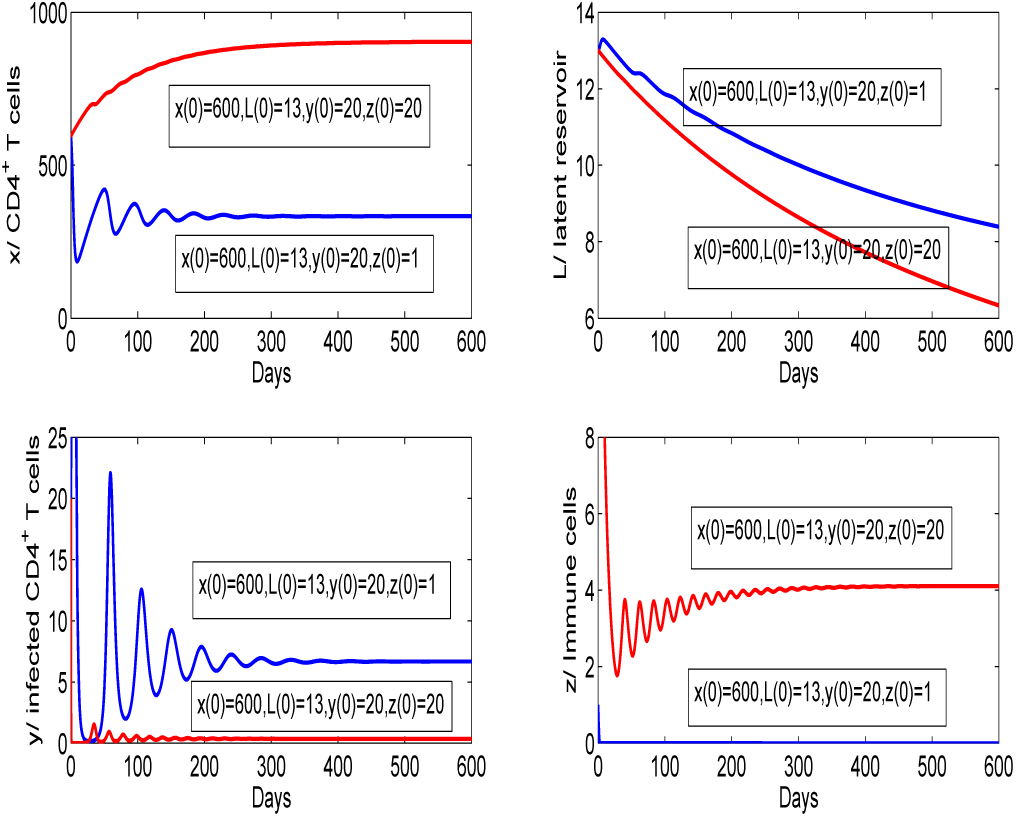
Time history of system (5.1). The trajectories of system (5.1) converge to different equilibria for different initial values, i.e., system (5.1) has bistable behavior. The initial values are *x* (0) = 600, *L*(0) = 13, *y* (0) = 20, *z* (0) = 1 (blue) and *x* (0) = 600, *L* (0) = 13, *y* (0) = 20, *z* (0) = 20 (red). The parameter values are shown in Table 5.

## Acknowledgments

This work was supported by the NSFC (No.U1604180) and Foundation of Educational Committee of Henan provence (No.19A110009).

